# Schizophrenia Risk Proteins ZNF804A and NT5C2 Interact at Synapses

**DOI:** 10.1101/2021.03.31.437821

**Authors:** Afra Aabdien, Laura Sichlinger, Nicholas J.F. Gatford, Pooja Raval, Madeleine R. Jones, Lloyd Tanangonan, Timothy R. Powell, Rodrigo R.R. Duarte, Deepak P. Srivastava

## Abstract

The zinc finger protein 804A (*ZNF804A*) and the 5′-nucleotidase cytosolic II (*NT5C2*) genes have been identified as robust susceptibility genes in large-scale genome-wide association studies of schizophrenia. The ZNF804A and NT5C2 proteins are highly expressed in developing and mature cortical neurons. ZNF804A has been implicated in regulating the development of neuronal morphology; it localises to synapses and is required for activity-dependent modifications of dendritic spines. NT5C2 has been shown to regulate 5′ adenosine monophosphate-activated protein kinase activity and implicated in influencing protein synthesis in neural progenitor cells. But despite these findings, a better understanding of the role these proteins play in regulating neuronal function is needed. A recent yeast two-hybrid screen has identified ZNF804A and NT5C2 as potential interacting proteins, but whether this occurs *in situ*; and moreover, in cortical neurons, is unknown. Here we show that ZNF804A and Nt5C2 colocalise and interact in hEK293T cells. Furthermore, their rodent homolouges, ZFP804A and NT5C2, specifically colocalise at synapses and form a protein complex in cortical neurons. Knockdown of *Zfp804A* or *Nt5c2* resulted in a significant decrease in synaptic expression of both proteins, suggesting that both proteins are required for the synaptic targeting of each other. Taken together, these data indicate that ZNF804A/ZFP804A and NT5C2 interact together in cortical neurons and indicate that these GWAS risk factors may function as a complex to regulate neuronal function.

## INTRODUCTION

Schizophrenia is a chronic psychiatric disorder of complex aetiology (Birnbaum and Weinberger, 2017; Lewis and Levitt, 2002; Rapoport et al., 2012; Smeland et al., 2020). Genome-wide association studies (GWAS) have suggested up to one third of genetic risk for schizophrenia arises from common alleles (Ripke et al., 2011) and the latest study has identified 270 independent schizophrenia risk loci (Ripke et al., 2020). The zinc finger protein 804A (*ZNF804A*) and the 5′-nucleotidase, cytosolic II (*NT5C2, cN-II*) genes were amongst the first ones to emerge from early GWAS of schizophrenia (O’Donovan et al., 2008 Ripke et al., 2011), and they have also been implicated in susceptibility to multiple psychiatric disorders (Amare et al., 2019; Cross-Disorder Group of the Psychiatric Genomics Consortium, 2013; Williams et al., 2011). Transcriptome-wide association studies (TWAS) have also associated *ZNF804A* (Gandal et al., 2018) and *NT5C2* (Hall et al., 2020) with schizophrenia. Ultimately, schizophrenia risk variants are associated with reduced expression of *NT5C2* (Duarte et al., 2016; Hall et al., 2020) and *ZNF804A* in the foetal and adult brain (Hill and Bray, 2012; Tao et al., 2014), but the function of these proteins in mature neurons remains unclear.

The ZNF804A protein has been found to be expressed in the human cerebral cortex, particularly in pyramidal neurons (Tao et al., 2014). ZNF804A has also been found to localise to putative synapses, as demonstrated by co-localisation with the synaptic proteins PSD-95 and GluN1 (Deans et al., 2017). Similarly, ZFP804A (rodent homologue) was found to be enriched in synaptic fractions, and to localise to dendritic spines, where the majority of excitatory synapses occur in the mammalian brain (Deans et al., 2017). Consistent with a role for the encoded zinc finger protein at synapses, knockdown or knockout of *Zfp804A* resulted in a reduction of dendritic spine density (Deans et al., 2017; Huang et al., 2020), whereas an increase in spine density was observed when the full length or shorter disease-associated isoform of the protein was overexpressed in cortical neurons (Dong et al., 2021; Zhou et al., 2020). Moreover, activity-dependent remodelling of dendritic spines was impaired in cortical neurons with reduced ZNF804A expression levels (Deans et al., 2017). Together, these data indicate a role for ZNF804A at synapses.

The NT5C2 protein also appears to be enriched in cortical neurons (Duarte et al., 2019), but relative to ZNF804A, less is known about the function of NT5C2 in the cortex. The knockdown of *NT5C2* in human neural progenitor cells is associated with increased phosphorylated 5′ adenosine monophosphate-activated protein kinase (AMPK), which negatively regulates mammalian target of rapamycin (mTOR) and eukaryotic elongation factor 2 (eEF2), suggesting a role in protein translation (Duarte et al., 2019). Knockdown of *CG32549* (the *Drosophila melanogaster* homologue) causes motor defects (Duarte et al., 2019), and increased sleep in flies (Singgih et al., 2021). Whereas it appears that *NT5C2* has a function in the developing and adult brain, the precise underlying neurological mechanisms are yet to be established.

A recent interactome study using a yeast two-hybrid assay has indicated that ZNF804A interacts with multiple proteins, including NT5C2 (Zhou et al., 2018). Follow up studies have demonstrated that ZNF804A forms a complex with several predicted interactors to regulate neuronal morphology (Dong et al., 2021). Interestingly, it was noted that NT5C2 is the only potential interacting protein for ZNF804A, encoded by a GWAS-supported gene (Zhou et al., 2018). In this study, we investigate the subcellular localisation of ZNF804A and NT5C2, and explored whether they formed a protein complex in heterologous hEK293T cells and rodent cortical neurons. We find that ZNF804A and NT5C2 do form a protein complex in heterologous and cortical neurons. Both proteins are present in multiple subcellular compartments but co-localise at or near synapses in cortical neurons. Small interfering RNA (siRNA)-mediated knockdown of either protein resulted in altered presence of both proteins at synapses. Together, these data support a model whereby two risk factors for schizophrenia may function as a complex to regulate synaptic function.

## RESULTS

### ZNF804A and NT5C2 form a protein complex in heterologous cells

As a yest-two hybrid assays has revealed that ZNF804A and NT5C2 are potential interacting partners (Zhou et al., 2018), we were interested in confirming this *in situ*. We first examined the localisation of ectopically expressed HA-ZNF804A and Myc-NT5C2 in hEK293T cells. Consistent with findings from Deans et al. (2017), HA-ZNF804A localised to both nuclear and cytosolic compartments **(Figure 1A)**. Within the cytosol and at the plasma membrane, HA-ZNF804A co-localised with the cytoskeletal filament, F-actin probe (phalloidin) **(Figure 1A)**. hEK293T cells transfected with Myc-NT5C2 showed that this protein mainly localised to the cytosol **(Figure 1B)**, consistent with findings from Duarte et al. (2019). We next co-transfected HA-ZNF804A and Myc-NT5C2 into hEK293T cells and observed that HA-ZNF804A and Myc-NT5C2 co-localised in the cytoplasm and near the plasma membrane of hEK295T cells **(Figure 1C)**. This was confirmed by a line graph construction, whereby the fluorescence intensity of both proteins was measured **(Figure 1D – i & ii)**, and the overlap between the intensities provided evidence of co-localisation.

**Figure 1.**
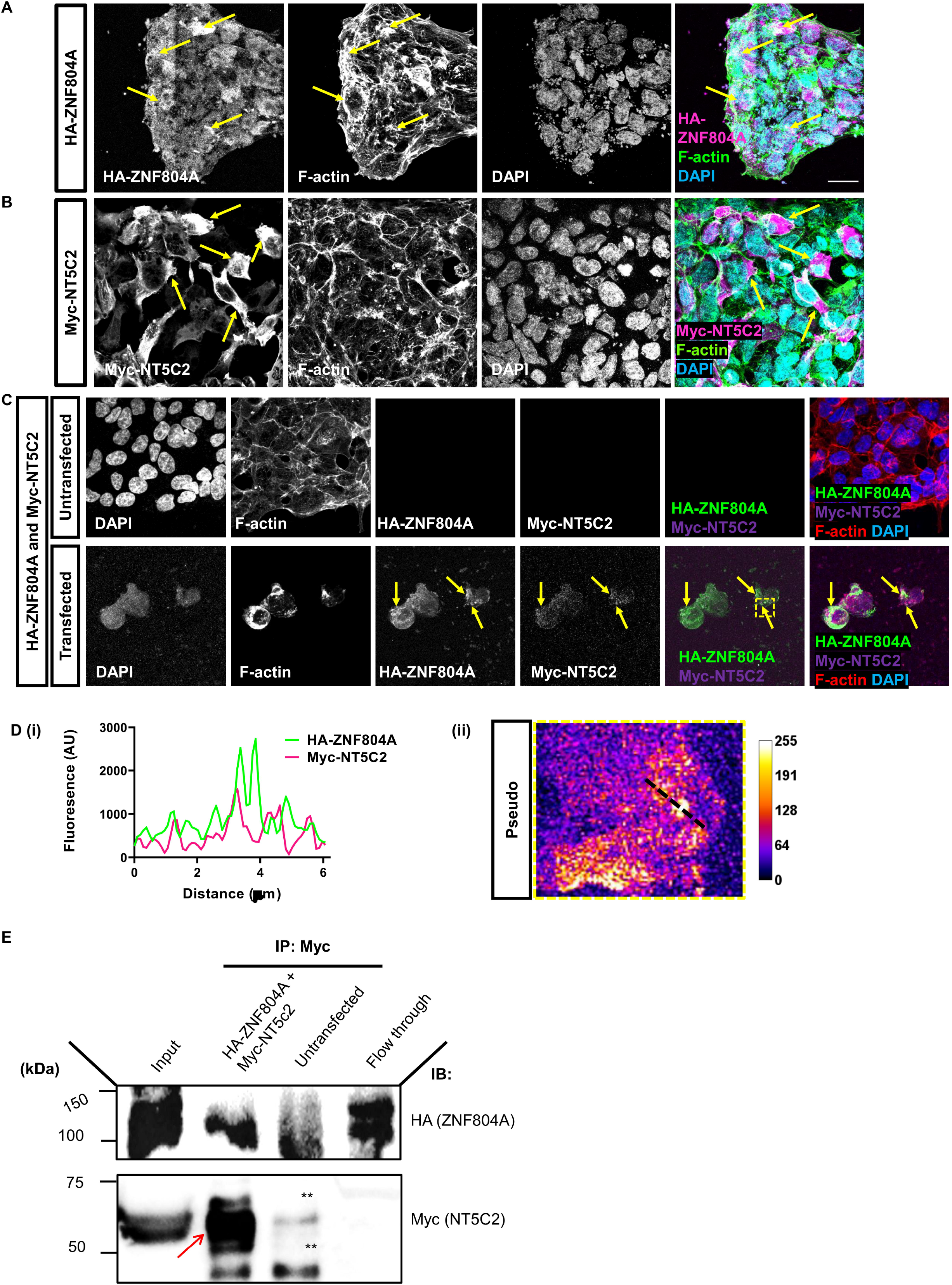
NT5C2 and ZNF804A form a protein complex and co-localise when expressed in hEK293 cells. **(A)** Representative confocal image of hEK293T cells expressing HA-ZNF804A and stained for F-actin. Ectopically expressed HA-ZNF804A localises to cell cytoplasm and plasma membrane (yellow arrows). Near the plasma membrane, HA-ZNF804A co-localises with F-actin. **(B)** Representative confocal image of hEK293T cells expressing myc-NT5C2 and stained for F-actin. Ectopically expressed myc-NT5C2 localises to cell cytoplasm (yellow arrows) similar to previous descriptions. **(C)** Confocal images of untransfected hEK293T cells and cells co-transfected with HA-ZNF804A and myc-NT5C2. In co-expressing cells, HA-ZNF804A and myc-NT5C2 are found to co-localise to the cell cytoplasm and near the plasma membrane (yellow arrows). Yellow dotted box indicates area magnified in (D). **(D) i) + ii)** Intensity plot and magnified image of dotted box from (C), of co-localised HA-ZNF804A and myc-NT5C2. **(E)** Co-immunoprecipitation (co-IP) assay of hEK293T cells co-expressing HA-ZNF804A and myc-NT5C2. Cell lysates were immunoprecipitated with a myc antibody to isolate NT5C2 and interacting partner. Samples were subjected to immunoblotting and then subsequently probed with antibodies for myc (to detect myc-NT5C2) and HA (to detect HA-ZNF804A). A specific band for HA-ZNF804A was detected in co-IP’ed cell lysates expressing both proteins but not from untransfected cell lysate. Scale bar = 5 µm.

To validate these findings and further determine whether ZNF804A and NT5C2 formed a protein complex, we performed a co-immunoprecipitation using hEK293T cells ectopically expressing both proteins. Cells co-transfected with HA-ZNF804A and Myc-NT5C2 were immunoprecipitated with an anti-Myc antibody to isolate NT5C2 and its interacting partners (**Figure 1E**). As expected, Myc-NT5C2 was readily detected in the immunoprecipitation (IP) sample. In addition, HA-ZNF804A was also present in the IP sample indicating that ZNF804A and NT5C2 were part of a protein complex **(Figure 1E)**. Taken together, these data indicate that ZNF804A and NT5C2 co-localise in the cell cytoplasm, near the membrane, and that they form a protein complex in hEK293T cells.

### Subcellular distribution of ZNF804A and NT5C2 in cortical neurons

Previous work has demonstrated that ZNF804A localises to somatodendritic compartments of neurons, and particularly in dendritic spines (Deans et al., 2017). In comparison, little is known about the distribution and organisation of NT5C2 in neurons. We first sought to confirm the ZNF804A distribution *in vivo*, using cell fractions generated from mouse cortex. Cortices were subjected to a subcellular fractionation protocol yielding whole cell (non-nuclear) (P1), cytosolic (S2) and crude synaptosomal (P2) subcellular fractions (Jones et al., 2014). Synaptosomal fractions were further subjected to Triton-X100 detergent separation to produce a supernatant (S) and precipitate (P) (Jones et al., 2014) **(Figure 2A)**. Consistent with our previous finding, ZFP804A was present in both cytosolic and synaptosomal fractions; however, the protein was highly enriched in synaptosomal supernatant, indicating a loose association with synaptic membrane **(Figure 2A)**. As expected, PSD-95, a major component of synapses and the post-synaptic density, was enriched in synaptosomal precipitate fractions indicating a tight association with synaptic membranes **(Figure 2A)**. Examination of NT5C2 distribution revealed that it was also present in cytosolic and synaptosomal fractions, although it was enriched in the cytosol. NT5C2 was only loosely associated with synaptic membrane as indicated by presence of the protein in synaptosomal supernatant fractions **(Figure 2A)**. These findings corroborate that ZFP804A /ZNF804A localises to multiple subcellular compartments beyond the nucleus (Deans et al., 2017), and suggest a role for NT5C2 at synapses.

**Figure 2.**
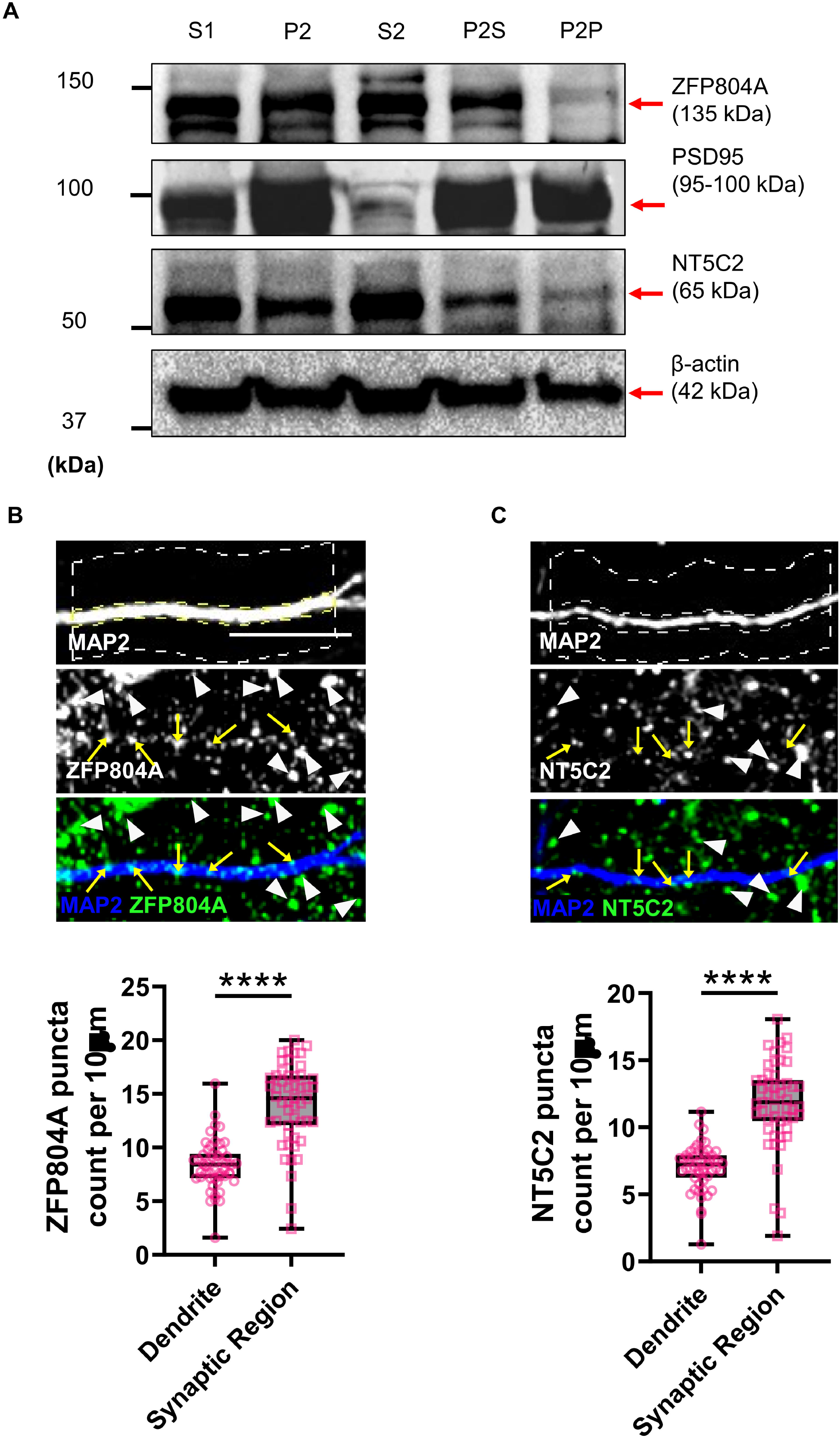
Synaptic localisation of ZNF804A and NT5C2 in cortical neurons. **(A)** Western Blotting of cell fractions generated from mouse cortex. S1, extranuclear cell lysate; P2, crude synaptosomal fraction; S2 cytosol; P2S, synaptosomal supernatant, P2P, synaptosomal precipitate. ZFP804A and NT5C2 were present in all fractions; ZFP804A and NT5C2 was enriched in P2S compared to P2P fractions, whereas NT5C2 was abundant in S2 fraction. PSD-95 was used to demonstrate synaptosomal enrichment, and β-actin as a loading control. **(B)** Representative confocal image of a section of dendrite from DIV20 cortical neurons immunostained for MAP2 (morphological marker) and a previously validated antibody for ZNF804A/ZFP804A. Yellow arrows indicate localisation of ZFP804A along dendrites, whereas white arrow heads indicate presence of protein in synaptic regions. Box and whisker plots, showing maximum and minimum values, is of ZFP804A linear density in either dendrites or synaptic region; n = 51 cells from 3 independent experiments. **(C)** Representative confocal image of a section of dendrite from DIV20 cortical neurons immunostained for MAP2 (morphological marker) and a previously validated antibody for NT5C2. Yellow arrows indicate localisation of NT5C2 along dendrites, whereas white arrow heads indicate presence of protein in synaptic regions. Box and whisker plots, showing maximum and minimum values, is of NT5C2 linear density in either dendrites or synaptic region; n = 51 cells from 3 independent experiments. Scale bar = 5 µm.

To further investigate the subcellular localisation of ZFP804A and NT5C2, we performed a series of immunostaining experiments in DIV20 (days *in vitro*) primary cortical neurons. Staining for endogenous ZFP804A revealed significant enrichment of this protein in the synaptic region (region adjacent to dendrite), compared to dendrites (ZFP804A puncta count: Dendrite, 8.37±0.32; Synaptic region, 14.03±0.53 (*t*(7)=9.10, *p*<0.0001, n=8)) **(Figure 2B)**. Staining for endogenous NT5C2 also revealed significant enrichment of this protein in the synaptic region compared to dendrites (NT5C2 puncta count: Dendrite, 7.01±0.24; Synaptic region, 11.77±0.45 (*U*=214, *p*<0.0001, n=8)) **(Figure 2C)**. Combined, these data indicate that ZFP804A and NT5C2 localise to both dendrites and synaptic regions, suggesting that both proteins are present within the same subcellular compartments in neurons.

### ZNF804A and NT5C2 colocalise and interact in cortical neurons

Considering that ZNF804A and NT5C2 form a protein complex in Hek293T cells, and that ZFP804A and NT5C2 localise to the same subcellular compartments in cortical neurons, we sought to explore whether these proteins were part of a protein complex in rat cortical neurons, *in situ*. First, we examined whether both proteins co-localised together in cortical neurons. In DIV20 cortical neurons, both ZFP804A and NT5C2 were found to co-localise in synaptic regions: this is demonstrated by the multiple white and yellow arrows and the green/magenta overlap in the orthogonal projections indicated by the orange arrows **(Figure 3A)**. To further confirm this synaptic colocalization, we analyzed ZFP804A puncta intensity within NT5C2 puncta in synaptic regions compared to dendrites. This revealed significantly more ZFP804A puncta overlapping within NT5C2 puncta in synaptic regions compared to dendrites (ZNF804A puncta intensity: Dendrite, 5.16±0.27; Synaptic region, 6.60±0.44 (*t*(7)=2.78, *p*=0.0065, n=8)) **(Figure 3B)**. The inverse relationship was also found to be true, whereby significantly more NT5C2 puncta were found to overlap with ZFP804A puncta in synaptic regions compared to dendrites (NT5C2 puncta intensity: Dendrite, 7.34±0.31; Synaptic region, 9.54±0.52 (*t*(7)=3.61, *p*=0.0005, n=8)) **(Figure 3C)**.

**Figure 3.**
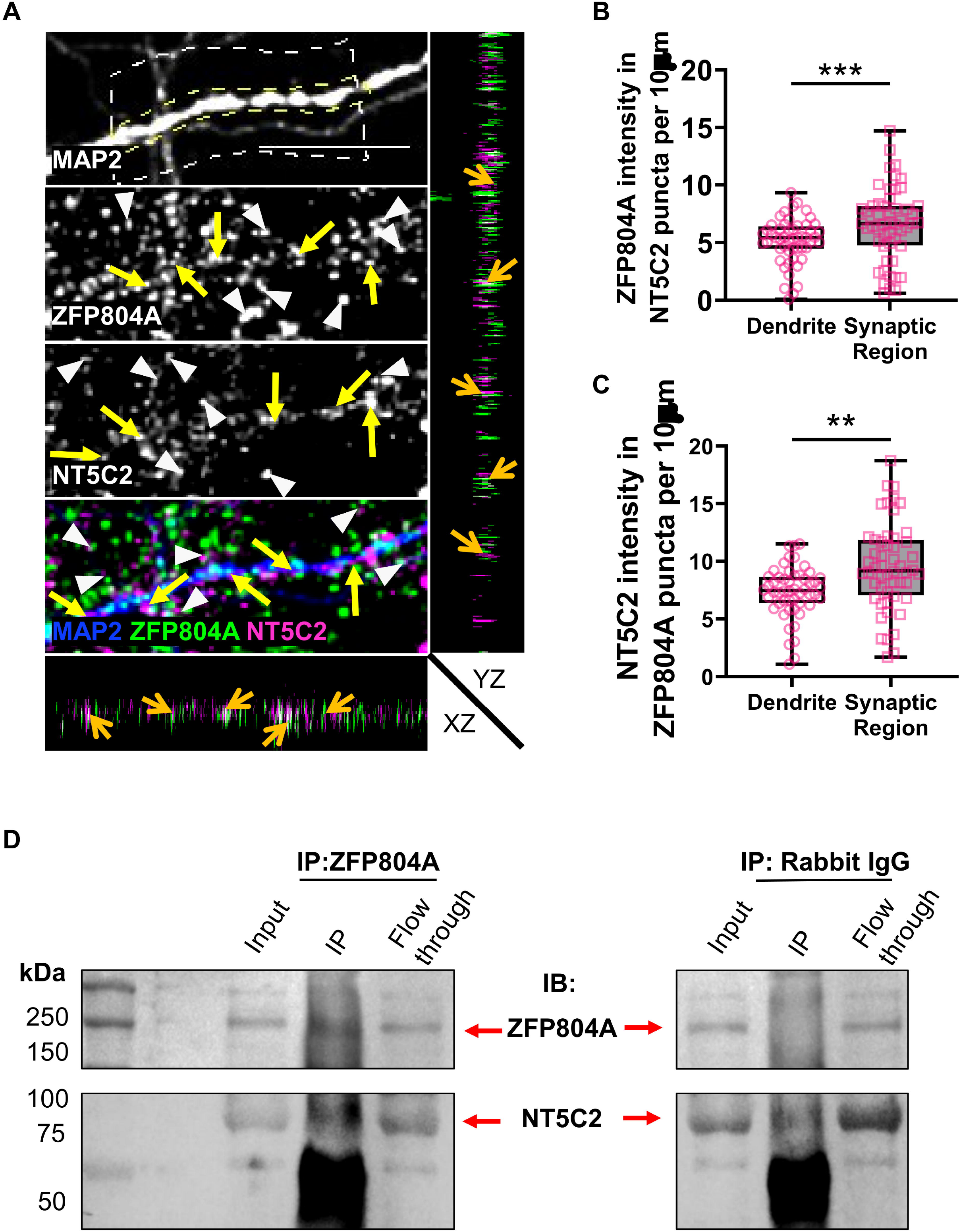
NT5C2 and ZNF804A co-localise and form a protein complex near synapses in cortical neurons. **(A)** Representative confocal image of DIV 20 cortical neurons immunostained for MAP2 (morphological marker), NT5C2 and ZFP804A. Yellow arrows indicate co-localisation of NT5C2 and ZNF804A along dendrites, whereas white arrow heads indicate co-localised puncta in synaptic regions. XZ and YZ orthogonal view further demonstrate co-localisation of both proteins (orange arrows). **(B and C)** Quantification of co-localisation: (B) density of NT5C2 puncta positive for ZFP804A and (C) density of ZFP804A puncta positive for NT5C2 staining along dendrites or within synaptic regions. N = 52 cells from 3 independent experiments. **(D)** Co-immunoprecipitation (co-IP) assay of DIV 20 cortical neurons. Cell lysates were immunoprecipitated with either an antibody against ZNF804A/ZFP804A or against rabbit IgG (control). Samples were subjected to immunoblotting and then subsequently probed with antibodies for NT5C2 and ZNF804A. A specific band for ZNF804A was detected in the input, ZNF804A-IP and flow through lanes as expected – no band was detected in the rabbit IgG-IP lane, indicating the specificity of the assay. Probing with an antibody against NT5C2 revealed bands in the input, ZNF804A IP and flow through lanes, but not the rabbit IgG-IP lane. This indicates that NT5C2 is part of a protein complex with ZNF804A. Scale bar = 5 µm.

To demonstrate this colocalization represented a potential interaction between the two proteins, we performed a co-immunoprecipitation assay for ZFP804A from cortical neuron lysates. Consistent with our data from heterologous cells, immunoblotting revealed that NT5C2 was present in lysates immunoprecipitated for ZFP804A, but not for rabbit IgG **(Figure 3D)**. These data indicate that ZFP804A and NT5C2 colocalize near synaptic regions in cortical neurons, and that they are part of a protein complex.

### Knockdown of *Zfp804a* or *Nt5c2* results in the redistribution of associated proteins

To further investigate a functional interaction between ZFP804A and NT5C2 in cortical neurons, we knocked down *Zfp804a* using a previously validated siRNA (Deans et al., 2017) in DIV15 primary rat cortical neurons. We then observed the effects of this knockdown on synaptic and dendritic NT5C2 localisation at DIV20. We first validated the efficacy of the *Zfp804a* siRNA by quantifying the number of ZFP804A puncta between *Zfp804a* siRNA knockdown, scrambled siRNA, or blank (untransfected) conditions. The *Zfp804a* siRNA was shown to successfully target ZFP804A as significantly fewer ZFP804A puncta were found in the siRNA knockdown condition compared to both the blank and scramble conditions (puncta per 10 µm: Blank, 22.45±1.44; Scramble, 22.05±1.03; *Zfp804a* siRNA, 16.38±1.20. One-way ANOVA: *F*(2,47)=27.04, *p*<0.0001, Blank/siRNA Bonferroni: (*t*(5)=6.91, *p*<0.0001, n=6); Scramble/siRNA Bonferroni: (*t*(5)=5.46, *p*<0.0001, n=6)) **(Figure 4A+B)**. No significant difference in ZFP804A puncta were observed between the blank or scramble conditions, indicating specificity of the siRNA treatment.

**Figure 4.**
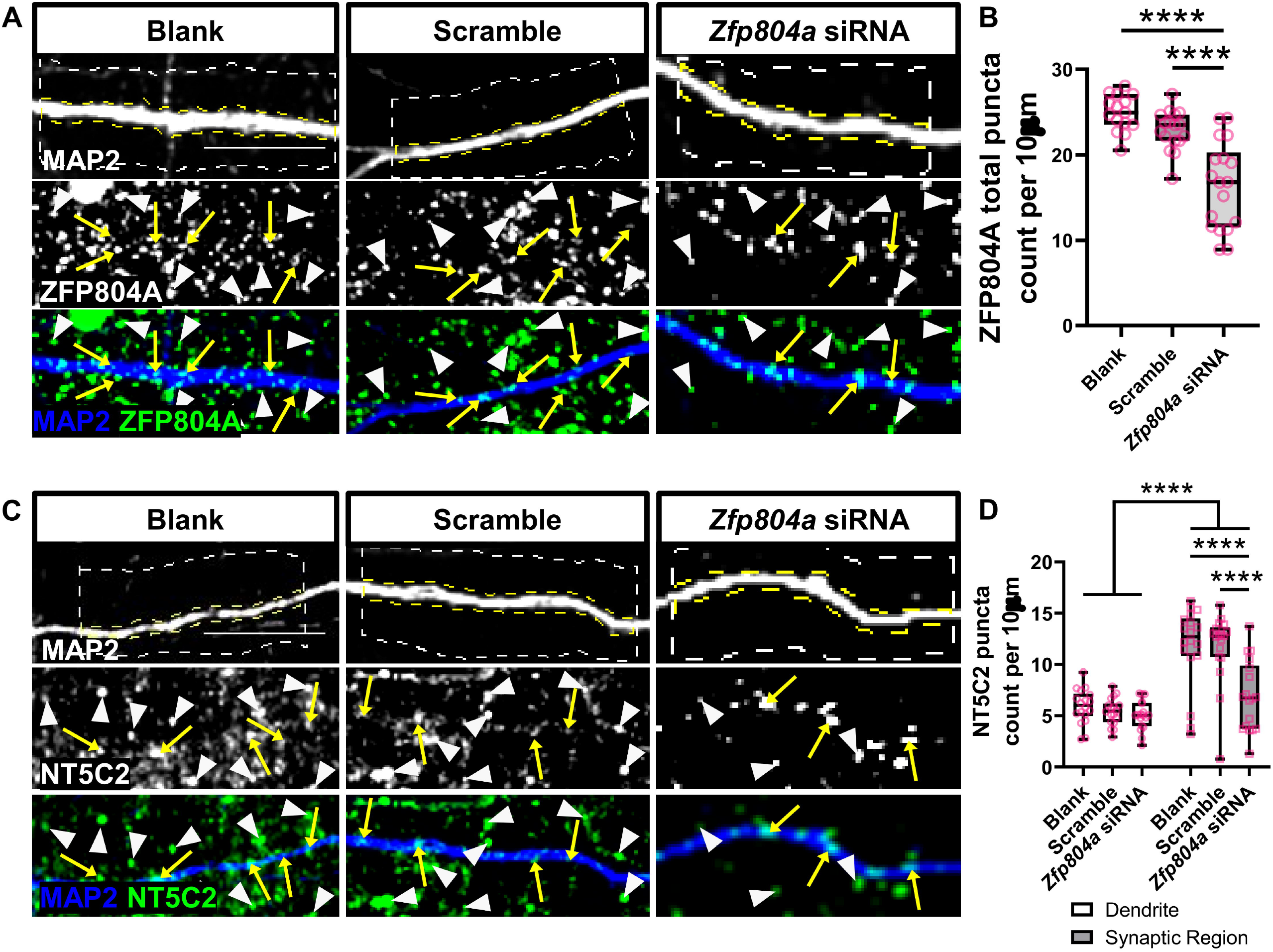
Knockdown of *Zfp804a* causes the redistribution of NT5C2. **(A)** Representative confocal image of DIV20 cortical neurons either untransfected (blank) or transfected with a scramble siRNA (scramble) or an siRNA for *Zfp804a* (*Zfp804a* siRNA). Cells were immunostained for MAP2 (morphological marker) or with an antibody against ZFP804A. Yellow arrows and white arrow heads indicate localisation of ZFP804A along dendrites or within the synaptic region respectively. **(B)** Quantification of ZNF804A linear density in Blank, scramble or *Zfp804a* siRNA conditions. Box and whisker plots show minimum and maximum values of linear density. N = 15-18 cells from 4 independent experiments. **(C)** Representative confocal image of DIV20 cortical neurons, transfected as in (A). Cells were immunostained for MAP2 and NT5C2. Yellow arrows and white arrow heads indicate localisation of NT5C2along dendrites or within the synaptic region respectively. **(D)** Quantification of NT5C2 linear density in Blank, scramble or *Zfp804a* siRNA conditions. Box and whisker plots show minimum and maximum values of linear density. N = 18 cells from 4 independent experiments. Scale bar = 5 µm.

Next, we knocked down *Zfp804a* in DIV15 rat primary cortical neurons and observed the effects on NT5C2 puncta in dendrites and synaptic regions of DIV20 cortical neurons. *Zfp804a* knockdown significantly reduced the density of NT5C2 puncta in synaptic regions (puncta per 10 µm; Synaptic region: Blank, 11.78±0.93; Scramble, 11.73±0.81; *Zfp804a* siRNA, 6.88±0.80. Two-way ANOVA: *F*(2,102)=7.29, *p*=0.0011; Bonferroni post hoc test (*p*<0.0001, n=6)) **(Figure 4C+D)**. In contrast, no effect on NT5C2 puncta number was observed in dendrites across conditions **(Figure 4D)**.

To further probe the nature of this functional interaction, we then performed the inverse experiment to understand if *Nt5c2* expression can influence ZFP804A localization. We knocked down *Nt5c2* using an siRNA previously validated (Duarte et al., 2019), in DIV15 rat primary cortical neurons. We then observed the effects of the knockdown on synaptic and dendritic ZFP804A localisation at DIV20, to allow for sufficient protein turnover. We first validated the efficacy of two siRNAs for *Nt5c2* via immunoblotting comparing between siRNA A or B knockdown, scrambled siRNA, or blank (untransfected) conditions. This revealed the *Nt5c2* siRNA A successfully targeted and knocked down NT5C2 in primary rat cortical neurons, as a clear decrease in protein expression was observed in this condition compared to both the siRNA B, blank, and scramble conditions **(Figure 5A)**. No difference in NT5C2 protein expression was observed between the blank or scramble conditions, indicating specificity of the siRNA treatment.

**Figure 5.**
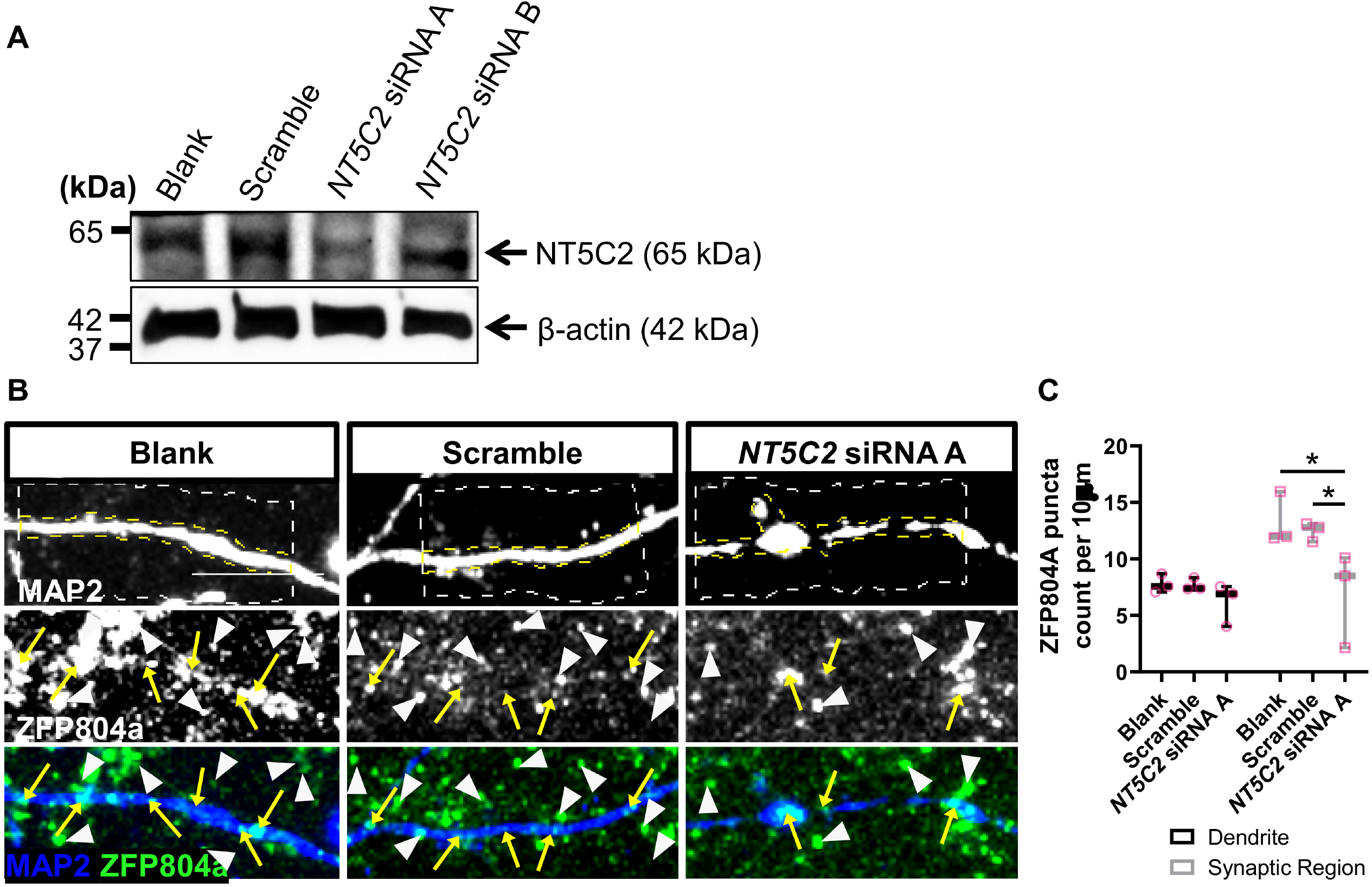
Knockdown of *Nt5c2* causes the redistribution of ZNF804A. **(A)** Western blotting of cell lysates from DIV20 cortical neurons either untransfected (blank) or transfected with a scramble siRNA (scramble) or an siRNAs (A or B) for *Nt5c2* (*Nt5c2* siRNA A or *Nt5c2* siRNA B). Immunoblots were probed with antibodies against NT5C2 or β-actin (loading control). **(B)** Representative confocal image of DIV20 cortical neurons, either untransfected (blank) or transfected with a scramble siRNA (scramble) or an siRNA against *Nt5c2* (*Nt5c2* siRNA A). Cells were immunostained for MAP2 and ZFP804A. Yellow arrows and white arrow heads indicate localisation of ZFP804A along dendrites or within the synaptic region respectively. **(C)** Quantification of ZNF804A linear density in Blank, scramble or *Nt5c2* siRNA A conditions. Box and whisker plots show minimum and maximum values of linear density. N = 15 cells from 3 independent experiments. Scale bar = 5 µm.

We knocked down *Nt5c2* in DIV15 rat primary cortical neurons and observed the effects on ZFP804A puncta in dendrites and synaptic regions of DIV20 rat primary cortical neurons. The knockdown significantly reduced the number of ZFP804A puncta in synaptic regions (puncta per 10 µm; Synaptic region: Blank, 13.28±1.34; Scramble, 12.47±0.48; *Nt5c2* siRNA A, 6.91±2.43. Two-way ANOVA: *F*(2,12)=6.07, *p*=0.0151; Bonferroni post hoc test (Blank/scramble, *p*=0.0113, n=3; Scramble/siRNA A, *p*=0.026, n=3)) **(Figure 5B+C)**. Similar to the ZFP804A knockdown, no effect on ZFP804A puncta was observed in dendrites when *Nt5c2* was knocked down. These data suggest a bidirectional signalling interaction occurs between NT5C2 and ZFP804A which operate in concert at synapses but not in dendrites.

## DISCUSSION

Mutations in multiple key synaptic genes have been previously associated with schizophrenia pathogenesis (Forrest et al., 2018; Penzes et al., 2011), but the precise nature of how these molecules interact to impair synapses in schizophrenia is unknown. *ZNF804A* and *NT5C2* are robust schizophrenia susceptibility genes (Ripke et al., 2020), that have been previously suggested to interact (Zhou et al., 2018). In this context, ZNF804A has been previously characterised as having roles in mediating activity-dependent structural plasticity of dendritic spines, neurite outgrowth, protein translation and gene transcription (Deans et al., 2017; Hill et al., 2012; Zhou et al., 2018). NT5C2, in turn, is a nucleotidase that regulates AMPK signalling, and has putative effects on protein translation (Duarte et al., 2019). However, the molecular mechanisms underlying their association with schizophrenia, or how they relate to synaptic function or to each other, remains unclear. Here, we show ZFP804A and NT5C2 colocalise and localise to synapses in cortical neurons, and that they are part of a protein regulatory network that is able to bidirectionally regulate itself, whereby expression of one protein regulates the expression of the other. This furthers our understanding of the role these proteins may play at synapses, and further suggest how both proteins may contribute to synaptic dysfunction in schizophrenia.

In this study we demonstrate that exogenous ZNF804A and NT5C2 form a protein complex in hEK293T cells. Further, we show that their rodent homologues, ZFP804A and NT5C2 are part of a protein complex in primary cortical neurons, although the consequence of this interaction is unclear. One biological function that both proteins appear to jointly regulate is protein translation. ZNF804A interacts with several proteins involved in protein translation, and has been shown to regulate the rate of protein synthesis in hEK293 and CAD cells (Zhou et al., 2018). NT5C2 has also been shown to regulate the activity of the 40S ribosomal protein RPS6, which is associated with protein translation (Duarte et al., 2019). Based on their predicted function and localization at synapses, we hypothesize that their interaction results in a protein complex (formed in combination with other proteins, as described in Zhou et al. (2018)) responsible for regulating local protein translation. Furthermore, defects in ZNF804A or NT5C2 may contribute to dysfunctional local synaptic protein synthesis in neurological disorders. Indeed, dysregulated protein synthesis has been shown in schizophrenia patient-derived olfactory neuronal cells and induced pluripotent stem cell-derived forebrain neural progenitor cells from schizophrenia patients (English et al., 2015; Topol et al., 2015). Common genetic variants in ZNF804A or NT5C2 could, therefore, be at least partly responsible for the dysregulation in protein translation observed in association with schizophrenia. Future studies are warranted to explore if a ZNF804A/NT5C2 protein complex could influence protein translation.

A second major finding in this study is the description of the subcellular distribution of ZNF804A and NT5C2 in cortical neurons. Both ZNF804A and NT5C2 were present in multiple subcellular compartments in cortical neurons. Consistent with our previous work (Deans et al., 2017), ZFP804A was present in cytosolic and synaptosomal fractions from cortical neurons. Interestingly, ZFP804A was predominately present in the supernatant fraction of synaptosomes following detergent separation. This indicates that ZFP804A is transiently, or ‘loosely’, associated with the synaptic membrane. NT5C2 was also present in both cytosolic and synaptosomal preparations; however, this protein was more abundant in the cytosolic fraction. In synaptosomal fractions, NT5C2 was also abundant in supernatant fraction of synaptosomes following detergent separation. The similar distributions of both proteins were further reflected by immunostaining. To note, our assessment of ‘synaptic regions’ by immunostaining as well as our subcellular fractionation protocol does not distinguish protein from being present in both pre- and post-synaptic compartments. Using super-resolution imaging, ZFP804A has been localised to presynaptic terminals as well as post-synaptic dendritic spines (Deans et al., 2017). It remains to be determined whether NT5C2 can be found in either, or both pre- and post-synaptic compartments. Nevertheless, our data indicates that rodent NT5C2 and ZNF804A colocalise in synaptic regions. Coupled with our co-immunoprecipitation data, this could indicate that these two proteins functionally interact in this region.

A limitation of our study is that, although the knockdown of either *Zfp804A* or *Nt5c2* in primary cortical neurons resulted in a loss of either protein specifically from synaptic regions, it is challenging to determine whether this was caused by a direct removal of the protein from synapses, or due to a loss of the synapses altogether. For example, siRNA-mediated knockdown of *Zfp804A* also results in the loss of dendritic spines (Deans et al., 2017). Therefore, the apparent reduction of NT5C2 in synaptic regions could be driven by the fact that there are fewer dendritic spines for the protein to localise to. NT5C2 can signal through AMPK; activation of this kinase has been shown to induce dendritic spine loss (Mairet-Coello et al., 2013). Thus, knockdown of *Nt5c2* could induce dendritic spine loss through AMPK activation. A better understanding of how synaptic structures are regulated will allow us to clarify the role of NT5C2 and ZNF804A proteins at synapses.

There is a growing appreciation of the need to understand how genetic risk factors interact with each other and with environmental factors. It is posited that such interactions likely shape and contribute to complex genetic disorders such as schizophrenia. Recent studies have begun to dissect how different genetic risk factors can act in a synergistic manner to converge on common biological processes to increase risk for psychiatric disorders (Schrode et al., 2019). In this study, we demonstrate that two genetic risk factors for psychiatric disorders co-localise and form a protein complex at synapses, thus revealing a novel understanding of these schizophrenia susceptibility genes. While additional research is required to understand the function of a ZNF804A and NT5C2 protein complex, these data support a model whereby genetic risk factors may directly work in a co-operative or non-additive manner to increase risk for schizophrenia.

## METHODS

### Cell Culture

Human embryonic kidney (HEK293) cells were maintained in a 37°C/5% CO_2_ atmosphere in DMEM:F12 (Sigma, D6421) supplemented with 10% foetal bovine serum (ClonTech, 631107), 1% L-glutamine (Sigma, G7513), and 1% penicillin/streptomycin (Life Technologies, 15070063). Cells were passaged and plated at 30-40% confluency on 1.5H 18 mm glass coverslips 24 hours prior to transfection.

Primary cortical neuronal cultures were harvested from pregnant Sprague-Dawley rats at embryonic day 18 as described previously (Srivastava et al., 2011). Animals were habituated for 3 days before experimental procedures, which were carried out in accordance with the Home Office Animals (Scientific procedures) Act, United Kingdom, 1986. Cells were seeded on 1.5H 13 mm glass coverslips coated with poly-D-lysine 10 μg/mL (Sigma, P0899) diluted in 1X borate buffer (Thermo Scientific, 28341) at a density of 3 × 10^5^ /well equating to 857/mm^2^. Cells were cultured in feeding media: neurobasal medium (21103049) supplemented with 2% B27 (17504044), 0.5 mM glutamine (25030024), and 1% penicillin/streptomycin (15070063) (all reagents from Life Technologies, UK). After 4 days in vitro (DIV), 200 μM of d,l-amino-phosphonovalerate (d,l-APV, ab120004; Abcam) was added to media to maintain neuronal health over long-term culture and to reduce cell death due to excitotoxicity (Srivastava et al., 2011). Fifty percent media changes were performed twice weekly until the desired time in culture was reached (DIV20), at which point cells were transfected or lysed.

### Transfection

hEK293T cells and primary cortical neurons were transfected using Lipofectamine 2000 (Life Technologies, 11668019). Briefly, 3 μl Lipofectamine 2000 was mixed with 1 μg of each HA-ZNF804A (Origene, RG211363) or Myc-NT5C2 (Origene, RC200194) plasmid construct in DMEM:F12 (Sigma, D6421) or neurobasal medium (primary cortical neurons only). DNA:Lipofectamine mixtures were incubated in a 37°C/5% CO_2_ atmosphere for 20 minutes before being added dropwise to plated hEK293T cells or primary cortical neurons. Transfected hEK293 cells were then further incubated in a 37°C/5% CO_2_ atmosphere for 24-48 hours before being fixed for immunocytochemistry or lysed for immunoblotting accordingly. Transfected primary cortical neurons were incubated in a 37°C/5% CO_2_ atmosphere overnight during transfection, then transferred back to feeding media the following day to maintain their viability before being fixed for immunocytochemistry or lysed for immunoblotting accordingly 24 hours later.

Short interfering RNA (siRNA) mediated knockdown of *Zfp804a* (the rat homolog of *ZNF804A*) in rat primary cortical neurons was conducted using the N-Ter Nanoparticle siRNA Transfection System (Sigma: N2788) at DIV15 per manufacturer’s instructions. The siRNA targeted exon 2 of *Zfp804a* and all results were compared to Blank (N-TER, no transfection), Scramble (negative control) conditions. Briefly, neurobasal feeding media was removed and replaced with neurobasal transfection media (containing all feeding media components, but without penicillin/streptomycin) with 1X APV prior to transfection. For 3 ml media, (1 ml per condition), the following concentrations were required: 14.04 μl of siRNA dilution buffer (Sigma: N0413) was added to 1 μl Double stranded deoxyribonucleic acid (dsDNA): Blank (N-TER), Scramble (negative control GC duplex, Thermo Scientific: 465372) or siRNA (Thermo Scientific: HSS150613). In a separate mixture, 7.2 μl N-TER was mixed with 37.8 μl nuclease free water. The transfection mixture and individual siRNA conditions were vortexed for 30 seconds and spun down with a microcentrifuge. Finally, 15 μl N-TER transfection mixture was added to each of the conditions, vortexed, spun down and kept in the dark for 20 minutes at room temperature. Each condition was added dropwise to the relevant coverslips and incubated at 37°C/5% CO_2_ for 5 days prior to immunocytochemistry.

siRNA mediated knockdown of *Nt5c2* in rat primary cortical neurons was conducted using the Trilencer 27-mer NT5C2 siRNA kit (Origene: SR307908) at DIV15 per manufacturer’s instructions. Briefly, primary cortical neurons were fed with 1X d,l -APV in neurobasal feeding media for 30 minutes prior to transfection. SiRNA A (sense sequence, 5′-UGAGAAGUAUGUAGUCAAAGAUGGA -3′) and siRNA B (sense sequence, 5′-ACAACUGUAAUAGCUAUUGGUCUTC -3′) conditions were compared to Blank (N-TER, no transfection), Scramble (negative control sense sequence, 5′-CGUUAAUCGCGUAUAAUACGCGUAT-3′). N-TER was mixed with Opti-MEM (Life Technologies: 31985-047) at a 1:50 ratio per manufacturer’s instructions. Mixtures were vortexed and spun down using a microcentrifuge, then kept in the dark for 30 minutes at room temperature. Each mixture was added dropwise to the relevant coverslips and incubated at 37°C/5% CO_2_ for 5 days prior to immunocytochemistry.

### Immunocytochemistry & Microscopy

Rat primary cortical neurons and hEK293T cells were fixed in 4% sucrose and 4% formaldehyde for 10 minutes with agitation. Coverslips were washed in phosphate buffered saline (PBS) and then incubated in ice cold methanol for 10 minutes. Cells were permeabilized in 0.1% Triton-X100 (Alfa Aesar, A16046) in PBS and blocked simultaneously in 2% normal goat serum (NGS – Cell Signaling Technology, 5425) in PBS for one hour with agitation. Coverslips were subsequently incubated with primary antibodies diluted in 2% NGS+PBS in a humidified chamber overnight at 4°C. The following day, coverslips were washed 3x in PBS for 15 minutes each and were incubated with secondary antibodies diluted in 2% NGS+PBS in a humidified chamber at room temperature for one hour. Coverslips were washed 3x in PBS for 15 minutes each. Lastly, cells were counterstained using 4’,6-Diamidino-2-Phenylindole (DAPI – Life Technologies, D1306) diluted in PBS before being mounted on microscope slides with ProLong Gold Antifade Mountant (Life Technologies, P36930) and left to dry for 48 hours prior to confocal imaging. Primary antibodies included anti-ZNF804A C2C3 rabbit polyclonal (GeneTex, GTX121178, 1:200), anti-NT5C2 mouse monoclonal (Abnova, H00022978-M02, 1:200), anti-NT5C2 rabbit polyclonal (Abcam, ab96084, 1:750), anti-MAP2 chicken polyclonal (Abcam, ab92434, 1:1000), anti-PSD95 guinea pig polyclonal (Synaptic Systems, 124014, 1:500), anti-GFP rabbit polyclonal (Origene, TA150032, 1:10,000), anti-HA mouse monoclonal (BioLegend, 901503, 1:1000), and anti-Myc mouse monoclonal (BioLegend, 626802, 1:1000). Actin filaments were stained using ActinGreen™ 488 (Life Technologies, R37110) or ActinRed™ 555 (Life Technologies, R37112) ReadyProbes™ per manufacturer’s instructions. Secondary antibodies included Alexa Fluor 488 goat anti-rabbit (A11034, 1:750), Alexa Fluor 568 goat anti-mouse (A11031, 1:750), Alexa Fluor 633 goat anti-guinea pig (A21105, 1:500), Alexa Fluor goat anti-chicken (A21449, 1:750) (all Invitrogen), and Alexa Fluor 405 goat anti-chicken (Abcam, ab175675, 1:500).

All experiments were imaged using a Leica SP-5 confocal microscope using a Plan-Apochromatic 63x 1.40 NA oil-immersion objective (Leica microsystems, 506210). Fluorophores were excited using a 100 mW Ar laser (458, 476, 488, 496, 514 nm lines), 10 mW Red He/Ne (633 nm), and 50 mW 405 nm diode laser. Z-stacks of each cell were acquired for all conditions at 0.5 µm step size using Leica Application Suite – Advanced Fluorescence software (LAS-AF; v2.7.3) installed on a corresponding desktop computer running Windows XP (Microsoft, SP3). All Z-stacks were exported to ImageJ (https://imagej.nih.gov/ij/v1.51j8) where maximum intensity projections and background subtracted images were generated (Schneider et al., 2012).

### Co-Immunoprecipitation, Cell Fractionation & Immunoblotting

hEK293 cells transfected with GFP-ZNF804A, HA-ZNF804A, or Myc-NT5C2 were lysed in co-immunoprecipitation (co-IP) buffer consisting of 10 mM Tris; pH7.4, 150 mM NaCl, 1% Triton-X100, 0.1% SDS, 1% Deoxycholate, 5 mM EDTA combined with a protease/phosphatase inhibitor cocktail consisting of 1 mM AEBSF, 10 µg/ml Leupeptin, 1 µg/ml Pepstatin A, 2.5 µg/ml Aprotinin, 0.5 M NaF, and 1% serine/threonine phosphatase inhibitor cocktail #3 (Sigma, P0044). Detergent soluble lysates were sonicated for 10 pulses at 35% power and centrifuged at 15,000 rpm for 15 minutes at 4°C to remove cell debris, samples were then placed on ice for 30 minutes. An input sample of 75 μl was removed from the lysate and protein concentration was assessed using a Pierce Bicinchoninic acid (BCA) protein assay kit (Thermo Scientific, 10678484). Concomitantly, 50 μl magnetic Dyna Beads Protein A (Invitrogen, 10002D) were washed in 500 μl tris buffered saline+0.05% NP40 wash buffer containing 8 μl ZNF804A antibody or 8 μl control Rabbit immunoglobulin (IgG) (Santa Cruz Biotechnology, sc-2027). Samples were incubated at 4°C for 3 hours on a rotator before being placed in a DynaMag™-2 magnetic rack (Thermo Scientific, 12321D) and washed twice with TBS+NP40 wash buffer. 150-200 μl total lysate was equally added to ZNF804A or control normal rabbit IgG-bound beads. The samples were then left rotating overnight at 4°C. The following day, co-IP samples were returned to the magnetic rack, 50 μl of flow through was removed, and underwent BCA analysis to determine protein concentration. The beads were washed in TBS+NP40 wash buffer and resuspended in 50 μl Laemmli sample buffer (Bio-Rad Laboratories, 161-0737) which were subsequently boiled at 95°C for 5 minutes to denature.

Cell fraction samples were prepared as follows. Cortical tissue from four 16 week old male CD1 mice were homogenized in 10x v/w of homogenisation buffer (0.32 M sucrose, 1 mM NaHCO_3_, 1 mM MgCl_2_) using a glass Dounce tissue homogenizer for 10 strokes. Samples were then centrifuged at 4°C and at 200 RCF for 5 minutes: the nuclear fraction (P1) was discarded, and supernatant containing the extranuclear cell fraction (S1) retained. An aliquot of the S1 fraction was then centrifuged at 14,000 RCF for 15 minutes to produce the cytosolic fraction (S2 -supernatant) and crude synaptosomes (P2 – pellet) fractions. The P2 pellet was resuspended in 1% Triton X-100 buffer (20% v/v Triton X-100) with inhibitors. A further aliquot from the resuspended P2 fractions were then centrifuged at 14,000 RCF for 15 minutes: supernatant contain the ‘lightly bound’ synaptic fraction (P2S); the pellet was resuspended in homogenisation buffer and contained the ‘tightly bound’ synaptic fraction (P2P). All samples were then stored at -80°C.

Samples were resolved by SDS-PAGE, transferred to a PVDF membrane and blocked for 1 hour in 5% bovine serum albumin (Sigma, A7906) in TBS+0.01% Tween20. Membranes were then immunoblotted with primary antibodies overnight at 4°C, followed by 3x 15 minute washes in TBS-T. Membranes were then incubated with anti-mouse (Life Technologies, A16078, 1:10,000) or anti-rabbit (Life Technologies, G-21234, 1:10,000) horseradish peroxidase (HRP) conjugated secondary antibodies for 1 hour at room temperature, followed by 3x 15 minute washes in TBS-T. Membranes were then incubated in Pierce electrochemiluminescence substrate (Thermo Scientific, 170-5061) for 5 minutes and subsequently scanned using a Bio-Rad ChemiDoc MP (Bio-Rad). Band intensity was quantified by densitometry using Image Lab software (Bio-Rad, v6.0.1).

### Quantification & Statistical Analysis

Dendritic spine puncta were quantified from secondary and tertiary dendrites of microtubule associated protein 2 (MAP2) positive neurons. Images of 3 neurons per condition per culture were analyzed of 6 biological replicate cultures, approximately 100 µm of the dendrite was selected from 2-3 different dendrites per image. The areas of the dendrite and spine region (2 µm on either side of the dendrite) were traced manually and thresholded to select puncta sizes of 0.08-2 µm^2^ for analysis. Puncta count within the dendritic region was compared to puncta count in the synaptic region in synaptic localization, synaptic colocalization and knockdown experiments. All images were background subtracted to exclude non-specific staining in ImageJ prior to quantification.

All datasets were subjected to outlier detection using the ROUT method (robust regression and outlier removal – (Motulsky and Brown, 2006)) in GraphPad Prism (v8.0.2 – Windows, GraphPad Software, San Diego, California USA,http://www.graphpad.com), detected outliers were removed from their corresponding dataset. All datasets were also tested for normality using the D’Agostino & Pearson normality test to identify which datasets required parametric or non-parametric analyses (D’agostino et al., 1990). All statistical analyses used an alpha level of 0.05. Non-parametric Mann-Whitney U test was used to analyse all synaptic localisation and synaptic colocalization data. Ordinary one-way analysis of variance (ANOVA) with Bonferroni correction for multiple comparisons was used for analysing ZFP804A puncta in ZFP804A knockdown experiments. Two-way ANOVA with Tukey’s test for multiple comparisons was used for analysing the effects of ZFP804A knockdown on NT5C2 puncta. Two-way ANOVA with Tukey’s test for multiple comparisons was also used for analysing the effects of NT5C2 knockdown on ZFP804A puncta. All data are shown as mean ± standard error of the mean (SEM) and all error bars represent SEM.

## Acknowledgements

This work was supported by grants from UK Medical Research Council, Grant No. MR/L021064/1 to DPS; UK Medical Research Council Centre for Neurodevelopmental Disorders (Grant No. MR/N026063/1); Royal Society UK (Grant RG130856) to DPS; Independent Researcher Award from the Brain and Behavior Foundation (formally National Alliance for Research on Schizophrenia and Depression (NARSAD) (Grant No. 25957), awarded to D.P.S.; L.S. is supported by the UK Medical Research Council (MR/N013700/1) and King’s College London member of the MRC Doctoral Training Partnership in Biomedical Sciences; P.R. was funded by a BBSRC-iCASE studentship (BB/M503356/1); R.R.R.D. received funds from the Coordenação de Aperfeiçoamento de Pessoal de Nível Superior (CAPES, BEX1279/13-0). We thank the Wohl Cellular Imaging Centre for their help with imaging.

## Author Contributions

A.A., L.S., N.J.F.G., P.R., M.R.J., L.T. and D.P.S. performed all experiments and subsequent analysis. D.P.S., T.R.P. and R.R.R.D. designed the project and experiments. A.A., N.J.F.G., and D.P.S. write the initial draft of the manuscript; A., L.S., N.J.F.G., P.R., M.R.J., L.T., T.R.P. and R.R.R.D. and D.P.S. edited and finalized the manuscript.

## Conflict of interest

The authors declare that they have no conflict of interest.

## Notes

### Competing Interest Statement

The authors have declared no competing interest.

